# Fundamentals of rapid injection molding for microfluidic cell-based assays

**DOI:** 10.1101/194605

**Authors:** Ulri N. Lee, Xiaojing Su, David J. Guckenberger, Ashley M. Dostie, Tianzi Zhang, Erwin Berthier, Ashleigh B. Theberge

**Author notes:** Corresponding author contact information: Ashleigh Theberge.

## Abstract

Microscale cell-based assays have demonstrated unique capabilities in reproducing important cellular behaviors for diagnostics and basic biological research. As these assays move beyond the prototyping stage and into biological and clinical research environments, there is a need to produce microscale culture platforms more rapidly, cost-effectively, and reproducibly. ‘Rapid’ injection molding is poised to meet this need as it enables some of the benefits of traditional high volume injection molding at a fraction of the cost. However, rapid injection molding has limitations due to the material and methods used for mold fabrication. Here, we characterize advantages and limitations of rapid injection molding for microfluidic device fabrication through measurement of key features for cell culture applications including channel geometry, feature consistency, floor thickness, and surface polishing. We demonstrate phase contrast and fluorescence imaging of cells grown in rapid injection molded devices and provide design recommendations to successfully utilize rapid injection molding methods for microscale cell-based assay development in academic laboratory settings.

## Introduction

Microscale cell-based and organotypic models have matured significantly over the last decade.^1,2,3,4,5,6^ As micro-technologies move beyond the proof-of-concept stage and into biological or clinical research, there is a critical need to produce larger quantities of devices with a higher level of reproducibility than is enabled by typical prototyping techniques.^4,7^ Here we demonstrate that ‘rapid injection molding’ is well poised to bridge the gap between proof-of-concept device development in bioengineering labs and the adoption by biological and clinical end users who require batches of hundreds to thousands of devices.

Traditional early-stage prototyping methods, such as computer numerical control (CNC) micromilling,^8^ hot embossing,^9,10,11,12^ soft lithography,^13,14^ and three-dimensional (3D) printing^15,16,17^ are inexpensive fabrication methods at low quantities with rapid iteration times during the device development phases. However, these methods typically require significant manual preparation steps (e.g. machine setup, master mold fabrication/PDMS preparation, device curing and cleaning) resulting in device-to-device variability and high device costs when part volumes are scaled up. Further, many protoyping techniques alter the materials or require specific materials (e.g., PDMS for soft-lithography and UV-curable resin for 3D printing) that can render the device incompatible with cell-based or other biology research applications.^18^, ^19^

Injection molding, on the other hand, is the gold standard for device manufacturing and enables high-throughput manufacturing of devices (in volumes of hundreds of thousands to millions) at low per-device costs, while maintaining tight tolerances and high reproducibility.^20,21^ Further, rapid injection molding enables device production in materials such as polystyrene (PS, typical material for cell cultureware),^18,22^ cyclic olefin copolymer (COC, superior optical properties for microscopy),^23,24^ and polypropylene (PP, resistant to organic solvents, a consideration for some sample preparation methods).

Traditional high volume injection molding approaches utilize complex molds fabricated using high quality steel with high precision milling. The tooling of these molds is usually expensive (sometimes exceeding $50,000) and induces significant lead times of up to 12 weeks. However, the molds are made to be highly durable and sustain millions of plastic injections. The high mold cost is typically out of budget for academic research projects, which seldom require millions of devices.

‘Rapid’ injection molding solutions have emerged in the last 10 years to facilitate the use of injection molding. Companies such as Proto Labs^®^(Maple Plain, MN, USA) and ARRK (San Diego, CA, USA) offer turn-around times of ∼2-15 days. Rapid injection molding sits at the intersection between prototyping methods and traditional high volume injection molding as it costs an order of magnitude less than high volume injection molding ($2,000 to $5,000 to produce a mold for rapid injection molding), allows the use of comparable materials and techniques, and offers rapid turn-around times.^25^ It is worth noting that rapid injection molding approaches have been proposed in academic settings using milled molds or nickel-plated wafers, often drawing on mold capabilities developed for hot embossing.^10,26,27,28^ These solutions may push the costs of the technique further down, though they require in-house expertise on these systems.

Rapid injection molding reduces the cost of traditional high volume injection molding by optimizing important steps in the process: (1) design verification and quoting is typically minimalistic and in certain cases is automated by proprietary computer software, (2) the molds are made of softer, less-durable, metals (e.g., aluminium or urethane) that can be produced faster and are only guaranteed for several thousand injections, and (3) the use of inserts that are much smaller and fit into a standard injection molding frame reduce the complexity of the mold tooling (also known as Master Unit Die - MUD Molds or quick change dies). Finally, rapid injection molding usually offers more limited surface polishing options that significantly add to the cost and therefore must be chosen sparingly to keep costs low. The shortcuts of rapid injection molding methods impose significant compromises on the fabrication capabilities and production quality. These can lead to potential sources of failure, particularly for the unique requirements of cell-based microscale systems.

Herein, we examine key features of importance for microfluidic device fabrication, with a focus on cell culture applications. We used 3D laser scanning confocal microscopy to quantify dimensions and image channel cross-sections across a range of aspect ratios in three materials (PS, COC, and PP). For cell culture applications, the roughness and thickness of the cell culture surface are important considerations for phase contrast and fluorescence microscopy. We tested the effect of polishing the metal mold on the surface roughness of the molded plastic; we probed the ability to make thin plastic microwells; and we tested the effects of polishing and plastic thickness on multicolor fluorescence microscopy. We show that injection molded channels can be solvent bonded to additional layers to create closed microchannels. Together this work highlights the versatility of rapid injection molding for open and closed microfluidic cell culture and imaging.

## Methods

### Device fabrication

Devices were designed using SolidWorks (2016); files for the devices are available in the ESI. Devices were manufactured using the Protomold service by Proto Labs^®^. The following methods were provided in brief from Proto Labs^®^. The device molds were micromilled from aluminum on a HAAS 3-axis CNC mill (HAAS Automation). The cost of the molds were as follows: the test device used in Figures 1-5 ($3,615) and the device for solvent bonding used in Figure 6 ($2,865). Devices were manufactured on an electric injection molding machine (Toshiba) using Styron^®^666D (PS; AmSty), Pro-fax 6323 (PP; M. Holland), and ZEONEX^®^480R polymer resin (COC; Zeon Chemicals). The test device (Figures 1-5) was manufactured in PS, PP, and COC from one mold, with the exception of those used in Figures S4 and S5. The closed cell culture device (Figure 6) was manufactured in PS. Surfaces on the molds for all devices were polished according to the schematic in Figure S10. PM-F0 (Blue): Non-cosmetic, finish to Proto Labs^®^discretion; PM-F1 (Gray): Low-cosmetic, most toolmarks removed; SPI-C1 (Yellow): 600 grit stone; SPI-B1 (Green): 600 grit paper; SPI-A2 (Red): Grade #2 diamond buff.

**Figure 1.**
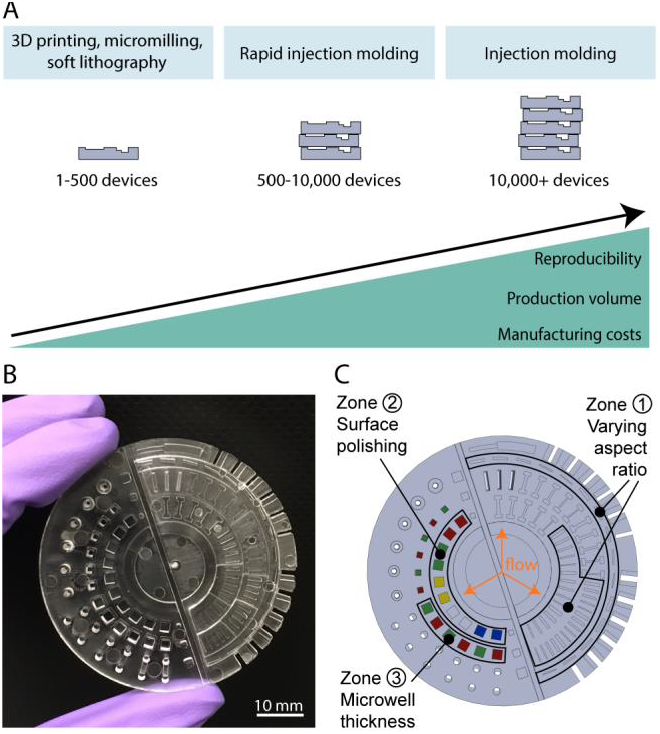
(A) Microscale device fabrication methods compared for their advantages and limitations. (B) Photograph of the PS device designed to test key characteristics of rapid injection molding for microscale cell-based applications. Photos of PP and COC test devices are shown in Figure S1. (C) Schematic of test device highlighting features of interest designed to test key requirements for microscale cell-based applications including rectangular channels positioned parallel and perpendicular to the plastic flow (zone 1), microwells for phase contrast microscopy (zone 2), and microwells for fluorescence microscopy (zone 3).

**Figure 2.**
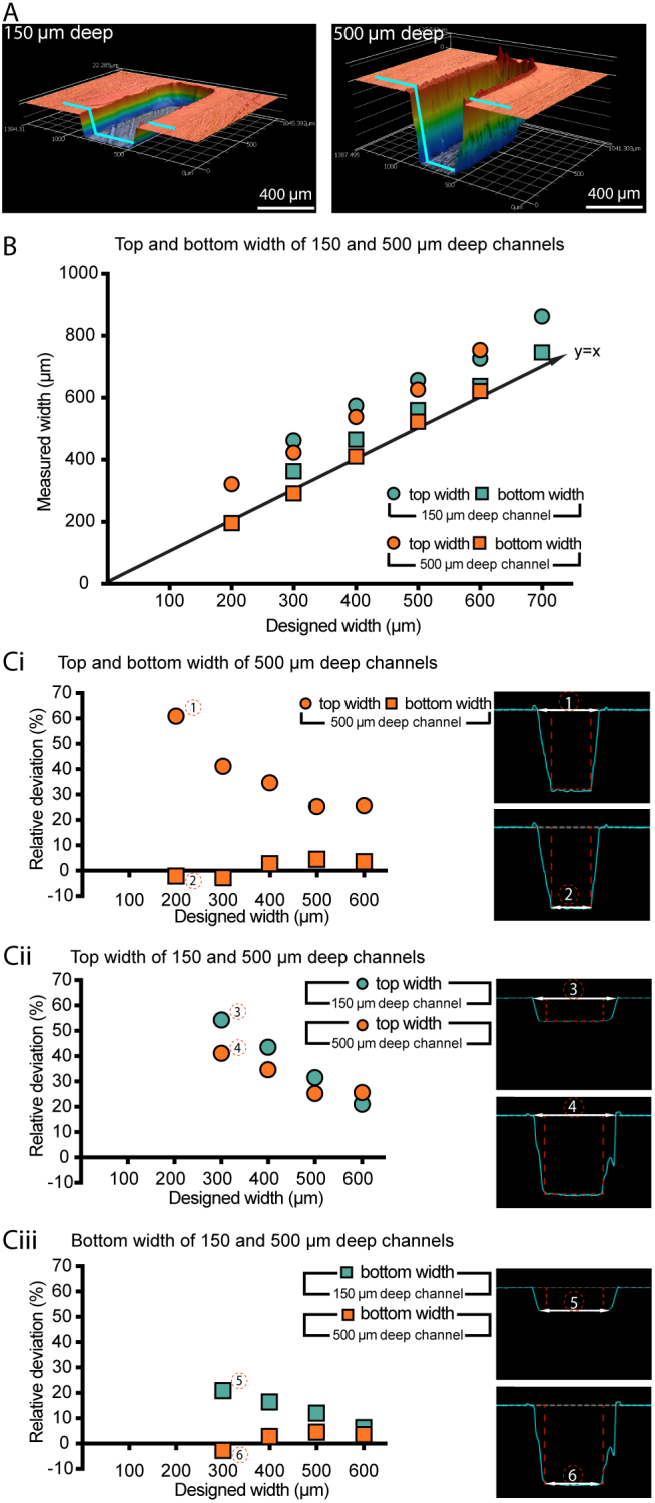
Quantification of dimensional differences between the device design and the injection molded device (PS device only). (A) Confocal microscope images of channels (designed width = 300 μm; left channel designed height = 150 μm, right channel designed height = 500 μm). The blue line indicates the cross section used for dimension measurements. Measurements of width were made at both the top and the bottom of the channel. (B) Comparison between the average measured width and the designed width for the 150 μm and 500 μm deep channels (the y=x line, representing ideal correspondence between designed and measured values). (Ci-Ciii) Plots of relative deviation for the parameters indicated (left) and corresponding confocal profilometer cross-sections (right). Data points represent the average of three devices produced by the same mold. In all cases, the standard error of the mean was smaller than the symbol plotted; the complete set of raw data is included in the SI, including separately plotted data points for the three replicate devices.

**Figure 3.**
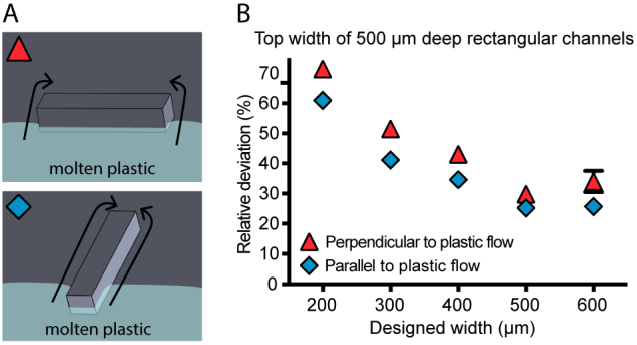
Effect of feature orientation relative to the flow of plastic in the mold for PS devices. (A) Schematic of molten plastic flow around mold for channels perpendicular (top) and parallel (bottom) to the flow of plastic. (Flow of plastic in the schematic is represented by light blue shading and arrows.) (B) Graph of average relative deviation in top width for rectangular channels parallel and perpendicular to the PS flow. Data points represent the average of three devices produced by the same mold. In cases where the standard error of the mean was smaller than the symbols plotted, error bars are not shown; the complete set of raw data is included in the SI, including separately plotted data points for the three replicate devices.

**Figure 4.**
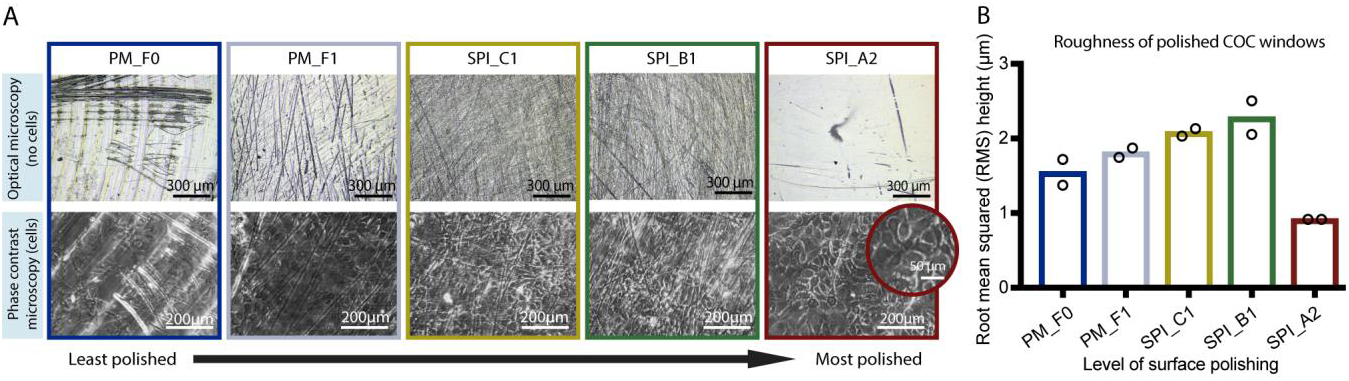
Effect of aluminium mold polishing on surface roughness of COC microwells and on clarity of phase contrast microscopy images of cells grown in COC microwells. The results indicate that the highest level of polishing (SPI_A2) is required for clear phase contrast images. Microwells corresponding to the data presented in this figure are shown in zone 2 of the test device schematic (Figure 1C), with the same color-coding to represent the level of polishing applied to the metal mold. Abbreviations “PM_F0” though “SPI_A2” correspond to the polishing level options offered by Proto Labs^®^ (see further explanation in Results section). (A) Top row: Optical microscopy images of COC microwells without cells imaged using a 3D laser scanning confocal microscope. Bottom row: Phase contrast microscopy images (10x magnification) of prostate epithelial cells (BHPrE1) grown on COC microwells; original images are included in Figure S9. Images are representative of two replicate microwells from one device. (B) Surface roughness (root mean squared (RMS) height) of COC surfaces measured using 3D laser scanning confocal microscopy. The bars indicate the mean of two replicate microwells from one device (data points from two replicates are superimposed on the bars).

**Figure 5.**
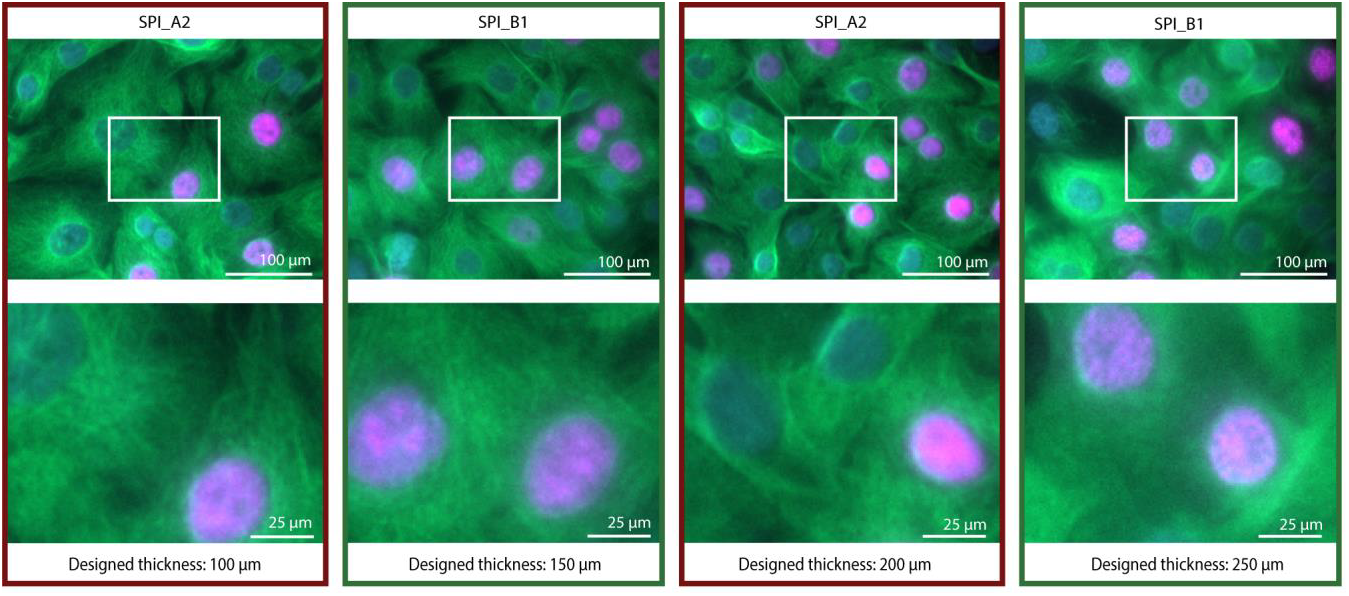
Fluorescence microscopy images of cells grown in COC microwells of varied thickness and polishing. Fluorescence images of prostate epithelial cells (BHPrE1) taken at 20x (0,40 NA) magnification with nuclear staining (DAPI, blue), proliferating nuclei (EdU, red), and tubulin (green). Zoomed in images of areas outlined in white are on the bottom row. The original images (with larger field of view) are included in Figure S7. The results indicate that the highest polishing level, SPI_A2, and a designed thickness of 100 μm enables clear images. Microwells corresponding to the data presented in this figure are shown in zone 3 of the test device schematic (Fig. 1C). The designed thickness is indicated in this figure; measurements of actual thickness are included in the SI, Figure S6A. Images bordered with red frames correspond to SPI_A2 polishing; green frames correspond to SPI_B1 polishing.

**Figure 6.**
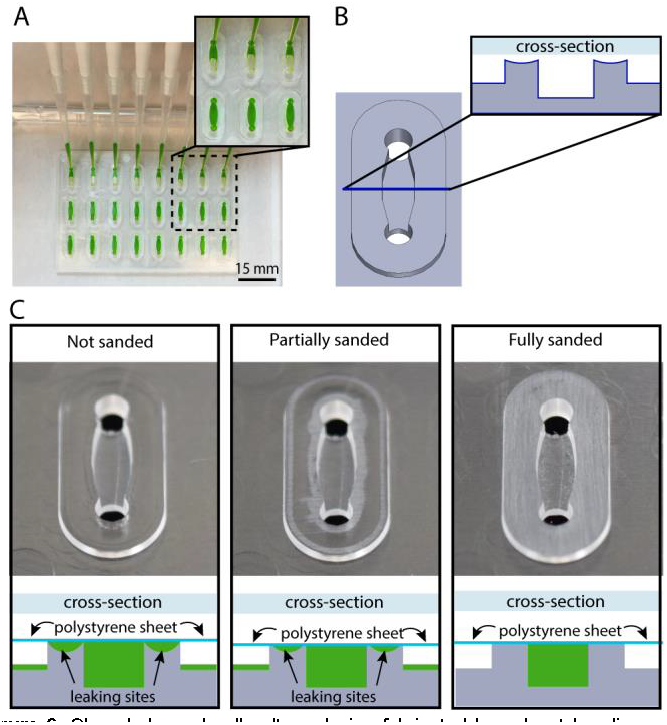
Closed channel cell culture device fabricated by solvent bonding a PS sheet (comprising the channel floor) and a PS injection molded device (comprising the channel walls and ceiling). (A) Closed channel cell culture device with multiplexed channel filling using a multichannel pipette. (B) Schematic of a single channel with oval island surrounding the channel. The cross-section (inset) shows that the oval island surrounding the channel is slightly curved due to plastic sinking caused by variable cooling rates of thick and thin regions of the injection molded device. (C) Schematic diagrams illustrating channel cross sections after solvent bonding to a PS sheet. For clarity the orientation of the schematic matches the orientation of the photographs directly above; when in operation, the device is flipped over such that the PS sheet forms the floor (as shown in A). The schematic diagrams indicate the results of filling the solvent bonded devices with fluid (green); leaking occurs when devices are assembled without sanding. Sanding provides a flat surface for solvent bonding enabling a leak-free device. Photographs show whitened areas of plastic removed by sanding the oval islands.

### Device cleaning

The devices were sonicated in 70% ethanol for 30 min using a M2800H ultrasonic cleaner (Branson). The devices were rinsed with fresh 70% ethanol and dried with compressed air. Note that this cleaning protocol is essential for cell viability; devices used directly from Proto Labs^®^without sonication resulted in poor cell viability as evaluated by cell morphology (likely due to the presence of mold release agents or other surface contaminates on the devices as received).

### Solvent bonding to create closed channel devices

The injection molded device, comprised of the channel walls and ceiling (shown in Figure 6), was sanded using 1200 grit sand paper. A 0.125 mm PS sheet (Goodfellow) was used for the channel floor. Both components were cleaned following the sonication protocol described above and dried with compressed air. Two cleanroom wipes (TX 609, TexWipe) were placed under the clean device on a programmable stirring hotplate (HS61, Torrey Pines Scientific) set to 37°C. The PS sheet was trimmed to cover the channels (∼ 45x70 mm). The PS sheet was set on the hotplate and acetonitrile (ACN) was deposited using a Pasteur pipette. The device was placed on top of the PS sheet with ACN. Excess ACN was removed using cleanroom wipes. A 310x77x6.35 mm aluminum bar (Alcoa) and 10 lb disc weight (CAP Barbell) were placed on top of the device for 5 min. The weight and aluminum bar were removed, and the solvent bonded device was left on the hotplate until the ACN fully evaporated (∼15 min). The device was transferred off the hot plate and placed between two room temperature aluminum bars for 20 min. The device along with aluminum bar and weight were placed back onto the hotplate, and the temperature was increased to 80°C. The device was heated for 2 h and left on the hotplate to cool overnight. For the image shown in Figure 6A,12.5 μL of 10% green food coloring (McCormick) was pipetted into each channel using an 8-channel multichannel pipette (Gilson).

### 3D laser scanning of rectangular channels and cell culture microwells

#### Device coating

Devices were coated with a gold/palladium (60:40) alloy for confocal profilometry and surface roughness measurements (Figures 2-4). The devices were sputtered with ∼16 nm of the alloy in a Quorum Q150R rotary-pumped sputter coater (Quorum Technologies) as follows: the pressure was reduced to 1e-1 mBar (3 min 43 s) and then to 6-8e-2 mBar (1 min); argon gas was released into the chamber (20 s) until the internal pressure reached 1 mBar; a gold/palladium alloy was sputtered at a current of 20 mA (2 min); the chamber was backfilled with argon gas until it reached atmospheric pressure. The process was repeated once more.

#### 3D laser scanning confocal microscope measurements with VK Viewer

Surface roughness and rectangular channel profile data were obtained using a 3D laser scanning confocal microscope (VK-X150, Keyence). The device was placed on a silicon wafer on the microscope stage. A 10X objective lens (Nikon) was used for all images. In the VK-Viewer software (Keyence) the Z-origin was reset before every scan, and expert mode was enabled. The zoom was set to 1.0X, the upper and lower position of the device were manually set for each scan, and no fixed Z-distance was set. Auto gain was enabled, and the neutral density (ND) filter was set to 10%. Brightness was not adjusted. Measurement settings were as follows: mode: surface profile, area: standard, and quality: high-accuracy. The real peak detection (RPD) was enabled, and the Z pitch was set to 3.00 μm.

#### Keyence image processing in MultiFileAnalyzer

MultiFileAnalyzer VK-H1XME v. 1.3 (Keyence) was used to analyze the images. In each 3D image representing a rectangular channel, the top surface on either side of the channel was fitted to a horizontal plane in order to remove natural tilt of the device. The profiles of the rectangular channels were taken and measured for top width, bottom width, and height using point-point analysis. The surface roughness of the square microwells was measured by normalizing the area of the microwell to a horizontal plane. The entire area of the scanned microwell was used for root mean square (RMS) height analyses.

### Microwell thickness measurements

The thickness of the microwells in the test device was measured using a 0-25 mm point and anvil micrometer (Fowler), with the micrometer tip placed in the center of the microwell.

### Plasma treatment

The devices were plasma treated using a Zepto plasma treater (Diener Electronic) for cell culture and liquid filling. (Devices used for 3D confocal imaging and surface thickness measurements were not plasma treated.) The chamber was pumped down to a pressure of 0.20 mbar, gas (air) was supplied (4 min, 0.25 mbar), and power enabled (2 min, 200 W).

### Cell culture microwells

#### Device preparation for fluorescence and phase contrast microscopy

After cleaning (see sonication protocol) and plasma treatment, devices were UV sterilized for 10 min in a Class II biosafety cabinet (Baker). 4 μL of 1% gelatin was pipetted into each microwell and incubated for at least 1 h at 37 °C. The gelatin was aspirated out of the microwell immediately prior to BHPrE1 cell seeding.

#### Maintenance of cells

BHPrE1 prostate epithelial cells^29^ were cultured in DMEM/F12 medium with 5% fetal bovine serum (FBS, VWR), 100 units/mL penicillin and 100 μg/mL streptomycin (Pen Strep, Gibco 15140122), 0.4% bovine pituitary extract (BPE, Hammond Cell Tech), 0.005 μg/mL epidermal growth factor (EGF, Sigma), and10 μg/mL insulin, 5.5 μg/mL transferrin, 6.7 ng/mL sodium selenite (purchased as 100x ITS, Gibco). Cells were used from passage 25-35. The cells were maintained in standard T25 culture flasks maintained at 37 °C with 5% carbon dioxide.

#### Cell culture in microwells

BHPrE1 cells were re-suspended at a density of 200 cells per μL. Then 4 μL of the suspended cells were seeded into each 2x2 mm microwell and cultured for 24 h. 5-ethynyl-2'-deoxyuridine (EdU, Invitrogen) was prepared according to the manufacturer’s specifications and diluted in cell culture media to 10 μM. 4 μL of the diluted EdU was added to each microwell. After 24 h, the cells were fixed with 4% paraformaldehyde (PFA).

#### Cell staining and imaging

The fixed cells were permeabilized with phosphate buffered saline (PBS) containing 0.5% Triton X-100 for 20 min and then washed twice with PBS containing 3% bovine serum albumin (BSA). Click-iT reaction cocktail (Invitrogen) was prepared following the manufacturer’s instructions and added to the cells, then incubated at room temperature for 30 min, during which the fluorophore (red, 647 nm) was attached. Cells were washed once with PBS containing 0.1% Triton X-100 (washing buffer), blocked with PBS containing 3% BSA for 1 h, and incubated with alpha tubulin monoclonal antibody (YL1/2, MA1-80017, Thermo Fisher Scientific) at 1:500 dilution overnight at 4°C. Cells were washed three times with washing buffer. The cells were incubated with goat anti-rat secondary antibody (1:200, green,488 nm, Jackson ImmunoResearch Laboratories) for 1 h at room temperature. The goat anti-rat secondary antibody was removed, a 1:2000 dilution of Hoechst (Invitrogen) was added, and the cells were incubated for 30 min. The cells were washed three times with washing buffer. Fluorescence images were taken using an Axiovert 200 inverted microscope (Zeiss) equipped with an AxioCam 503 mono camera using a 20x (0.40 NA) objective. Phase contrast images were taken on a Primovert microscope (Zeiss) with a MU1403B camera (Amscope). Brightness/contrast adjustments were made uniformly across all images, and the original unadjusted files are included in Figures S7 and S9.

## Results and Discussion

Rapid injection molding is a technique that is gaining popularity as it sits at the intersection of prototyping and traditional high volume injection molding. It enables the production of ∼500-10,000 devices while keeping setup costs low and providing little inter-device variability (Figure 1A). These features make it an ideal option for the fabrication of microscale devices during the transfer from design prototyping to initial applications in biology research laboratories or for clinical studies.

In general, injection molding is a process where by molten plastic is injected into a cavity formed by two (typically metal)molds; the plastic is allowed to cool, then ejected as a solid part (i.e., device). In the context of microfluidic devices, it can be difficult to maintain fidelity in fine features (particularly features with dimensions <0.5 mm). Rapid injection molding companies often utilize proprietary software to screen a customer’s designs and flag features that are known to impact feature fidelity, such as the radii of curvature of edges, excessively thick/thin regions, or improper drafting. However, while the design criteria are well known and understood for large parts and macroscale features, we have found that many of the features flagged by the software can actually be fabricated. Therefore, we sought to systematically determine the limits of rapid injection molding for microscale systems and develop strategies to overcome these limitations.

To test the limits of rapid injection molding for microfluidic devices we designed a test device (Figure 1B and 1C) that contains typical features of microfluidic devices, such as channels of various aspect ratios (Figure 1C, zone 1). Further, we designed areas of the test device to simulate cell culture surfaces to assess (1) the impact of different polishing options on the ability to image cells using phase contrast microscopy (Figure 1C, zone 2) and (2) the ability to create thin microwells for fluorescence microscopy applications (Figure 1C, zone 3).

### Effect of channel aspect ratio and orientation on deviation from designed dimensions

The channels included on the test device represent typical channel geometries commonly used in microfluidic cell culture applications. We designed channels with rectangular cross-sections of varying aspect ratios; the channel depths ranged from 150 μm to 500 μm, and the widths ranged from 200 μm to 700 μm. The channels were imaged using a confocal profilometer, and a cross-section of the 3D image (Figure 2A) was taken to measure the bottom width, top width, and height. Each of these measurements were compared to the designed dimensions in the CAD model. The devices were molded in polystyrene (PS), cyclic olefin copolymer (COC), and polypropylene (PP) – materials commonly used for cell culture and biological applications. It is worth noting that we used the same mold for all three materials (to reduce costs), and that the mold/parameters were optimized by Proto Labs^®^to accommodate the thermal shrinkage of PS. Since PP and COC have different thermal shrinkage rates than PS, we would expect deviations in the dimensions across the materials. Thus, data for PP and COC are provided in the Supplementary Information (Figures S2 and S3) as a reference for the reader; we caution against strict comparisons of the dimensions between the three materials.

We present here the results from the PS device (Figure 2) as well as measurements collected from two different batches of PS devices, prepared on two separate metal molds, both with the same design (Figures S4 and S5). We find that the bottom widths are smaller than the top widths for all channel heights (Figure 2B) despite the fact that no drafting was used (i.e., the walls of the channel should be vertical). The difference in widths is approximately constant (in absolute value) across all channel widths and heights. The linear offset is a possible result of tolerancing issues during the machining of the mold, the wear on the milling tools used to create the mold, or polishing during post-processing (Figure 2B). As shown in Figure 2B, the average measured bottom widths in the 500 μm deep channels were smaller than the corresponding average measured top widths by approximately 90-130 μm. Additionally, the linear absolute change means small features will have larger relative percent deviation from design (Figure 2Ci). The relative difference in widths was also larger for shallower channels (i.e., 150 μm) compared to deeper channels (i.e. 500 μm, Figure 2Cii-Ciii). Similar patterns were found for COC and PP (Figures S2 and S3).

The device was designed with a circular layout with the gate (i.e., the injection location of molten plastic) at the center in order to obtain a radial flow of plastic with a controlled direction and even equi-radial velocity (Figure S10A). As expected, our results show that the direction the molten plastic fills the mold is relevant when designing a microfluidic device for injection molding. Rectangular channels were oriented either perpendicular or parallel to the flow of plastic to determine if the direction of molten plastic flow around the features affects the fidelity of the channels (Figure 3A). The channels perpendicular to the flow of plastic deviate more in top width than the channels parallel to the flow of plastic (Figure 3B). For the perpendicular channels, the increase in deviation from the designed width could be a result of the molten plastic improperly filling behind the metal feature as it flows over it, compared to easily flowing on each side of a metal structure placed to the direction of the flow (Figure 3A). This observation indicates a potential benefit of placing the gate of injection molded microfluidic device in the main axis of the channels to prevent a perpendicular flow of molten plastic across a channel. We present here the results from the PS device (Figure 3), and the equivalent information collected on COC and PP is included in the Supplementary Information (Figure S2 and S3) as well as measurements collected from two different batches of PS devices, prepared on two separate metal molds (Figure S4 and S5).

### Optical microscopy and surface roughness

Optical microscopy is a widely used technique for imaging cell morphology that requires good optical properties of the microfluidic materials. The aluminium mold from which the plastic devices were produced was fabricated using CNC milling, which inherently leaves tooling marks in the metal mold. We endeavored to determine the level of mold polishing required to enable optical microscopy. Polishing the mold adds to the cost, with higher costs associated with higher polishing levels, so it is important to determine the minimum polishing that enables satisfactory imaging. We studied five levels of polishing offered by Proto Labs^®^: PM-F0: Non-cosmetic, finish to Proto Labs^®^discretion; PM-F1: Low-cosmetic, most toolmarks removed; SPIC1: 600 grit stone; SPI-B1: 600 grit paper; SPI-A2: Grade #2 diamond buff (see Figure 1C, zone 2 of the test device).

We measured roughness using the root mean squared (RMS) height of the tooling and polishing marks on the microwell surfaces with 3D laser scanning confocal profilometry (Figure 4). The results indicate that the RMS increased with polishing from PM-F1 to SPI-B1 (Figure 4B), and the images (Figure 4A, top) show that from PM-F1 to SPI-B1 increased polishing only served to remove large tooling marks but increased smaller scale scratches on the device surface. The RMS decreased only at the highest level of polishing, SPI-A2 (Figures 4A, top and 4B). Correspondingly, only SPI-A2 polishing enabled sufficient clarity for phase contrast microscopy imaging of prostate epithelial cells (BHPrE1 cells) (Figure 4A, bottom). The data and images shown in Figure 4 correspond to the COC device. Surface roughness measurements for PS, another optically viable material for microscopy and common cell cultureware material, showed similar results to COC (Figure S8).

### Fluorescence microscopy

Fluorescence microscopy is a ubiquitous and quantitative tool used in cell-biology research that often requires high-magnification imaging to observe sub-cellular features. In addition to polishing that affects the ability to resolve cellular features as described previously, the thickness of the plastic can affect the resolution of imaging (some microscope objectives are tuned to be used with cover slips of specific thickness with a specific refraction index). In our test device, we included microwells with plastic thicknesses ranging from 100 μm to 350μm, increasing in 50 μm increments, and with alternating polishing levels between SPI_B1 and SPI_A2, the two highest levels of polishing available at Proto Labs^®^(see Figure 1C, zone 3 of the test device). These dimensions span typical thicknesses of glass cover slips. We measured the actual thickness of the floor of the COC microwells to be ∼40-50 μm greater than designed (Figure S6A). The PS devices were ∼5-30 μm greater than designed (Figure S6B) and the PP devices were ∼0-20 μm greater than designed (Figure S6C). Thus, the COC microwell with a designed thickness of 100 μm resulted in an actual thickness of 150.4 ± 2.4 μm, corresponding closely to No. 1.5 glass coverslips (thickness of 160-190 μm), commonly used in microscopy. While microwells of 100 μm designed thickness formed in all COC and PP devices (n=50), the microwells did not form in PS devices (these molded as undesired through holes in the device) (Figure S6B). Creating thin plastic features of dimensions compatible with high-magnification microscopy is thus possible using rapid-injection molding. Further, our results indicated the highest Proto Labs^®^polishing level (SPI_A2, Grade #2 Diamond Buff) on the thinnest COC microwells (i.e., designed thickness 100 μm, actual thickness 150.4 ± 2.4 μm) enabled multicolor fluorescence microscopy. An example image showing clarity in the tubulin immunofluorescence stain and nuclear stains (DAPI and EdU) in BHPrE1 prostate epithelial cells is shown in Figure 5.

### Fabricating closed channels with solvent bonding

The creation of microfluidic devices from injection molded devices often requires the bonding of an additional layer of plastic to create closed cavities.^23,30,31^ Figure 6A shows an example of a polystyrene cell culture device in which the channel walls and ceiling were produced by injection molding and the floor of the channel is a polystyrene sheet that is bonded to the injection molded device using acetonitrile solvent bonding. The device is based on a design previously published for primary testis cell culture studies.^32^ Maintaining a planar surface to allow reliable solvent bonding presents design challenges in the context of injection molding. Areas of varying thickness result in regions of the device cooling and solidifying at different rates and thus variable shrinking arises. Thicker regions of the device can lead to a phenomenon called ‘sinking’ in which visible deformation of the plastic surface occurs. In the design presented, the channels were 1.0 mm in depth. As that thickness across the whole device would have led to sinking, the device was designed as an array of 1.5 mm thick oval ‘islands’ in which the channels were molded (Figure 6, design file provided in SI). The space in between these raised island structures also provided an area for solvent to evacuate during the solvent bonding process.

In practice, we found that the cross sections of the oval islands surrounding the channel were slightly curved (Figure 6B); potentially due to the uneven cooling rates of the thicker plastic in the island regions compared to the thinner plastic on the remainder of the device. The curvature of the island prevents the polystyrene sheet from properly bonding to the device and results in channel leakage (shown schematically in Figure 6C, bottom). To solve this problem, we manually sanded the surface of the device to remove the top layer of plastic until it was flat, enabling successful solvent bonding (Figure 6C, top) and a leak-free array of closed channel devices (Figure 6A).

## Conclusions

We demonstrate here that rapid injection molding is an efficient and attractive technique to bring microscale platforms from the initial design stage to a reproducible, medium volume, application phase. Rapid injection molding enables the production of microdevices in volumes of 500 – 10,000 at mold costs of $2,000-5,000 with little inter-device variability. Importantly, injection molding enables the selection of materials that have demonstrated advantages for cell culture (e.g., polystyrene) and optical properties (e.g., COC). We show that rapid injection molding induces a small amount of deformation to the rectangular cross-section of a channel, though channels with a width of as low as 200 μm were produced with good precision. Further, we show that rapid injection molding can produce chambers for cell culture that are compatible with optical microscopy and fluorescence microscopy – an essential feature for cell-based microfluidics and organotypic microscale models. Finally, rapid injection molding is compatible with microdevice construction using solvent bonding methods to create closed channels. Further, rapid injection molding is optimally suited for capillary-driven open microfluidic techniques in which the devices can be utilized directly after injection molding, without further fabrication steps.^33,34^ With the right design considerations, rapid injection molding is a technique that is compatible with academic resources and that has the potential to enable the translation of technologies toward biological or clinical research.

## Acknowledgements

This work was supported by NIH K12 DK100022, the University of Washington, and through an award from the Kavli Microbiome Ideas Challenge, a project led by the American Society for Microbiology in partnership with the American Chemical Society and the American Physical Society and supported by The Kavli Foundation. We thank Prof. Simon Hayward for providing the BHPrE1 prostate epithelial cells and Dr. Mark Morgan for providing training and helpful discussions on the instrumentation used at the Washington Nanofabrication Facility.

## Notes and references

§ The authors acknowledge the following potential conflicts of interest: EB: Tasso, Inc., Salus Discovery, LLC and Stacks to the Future, LLC, DJG: Salus Discovery, LLC and Tasso, Inc., ABT: Stacks to the Future, LLC.

---

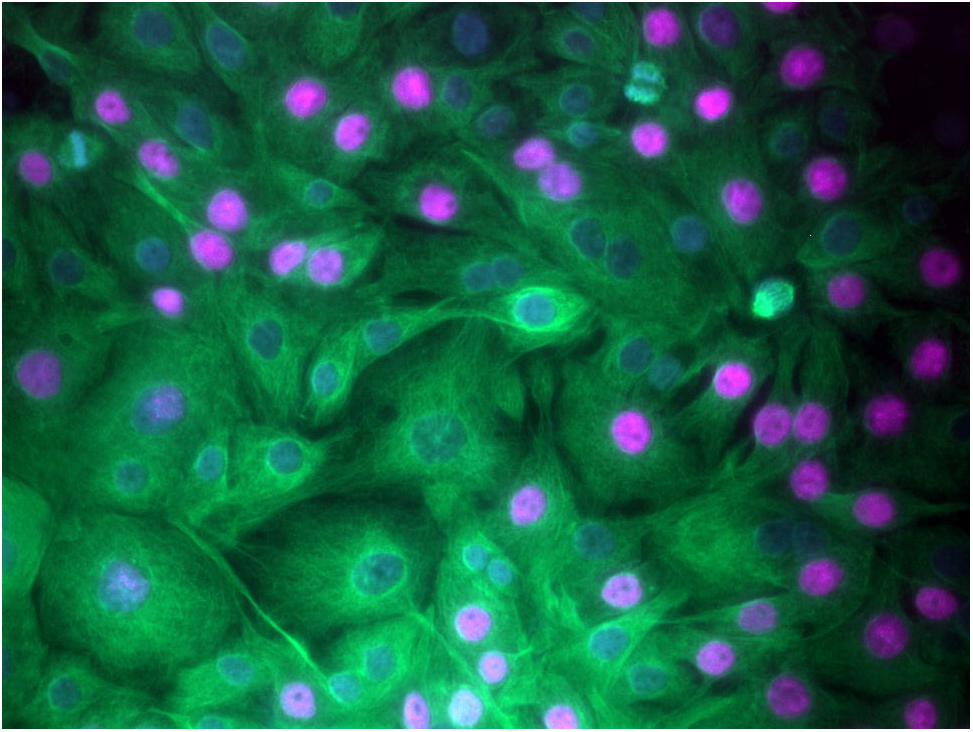

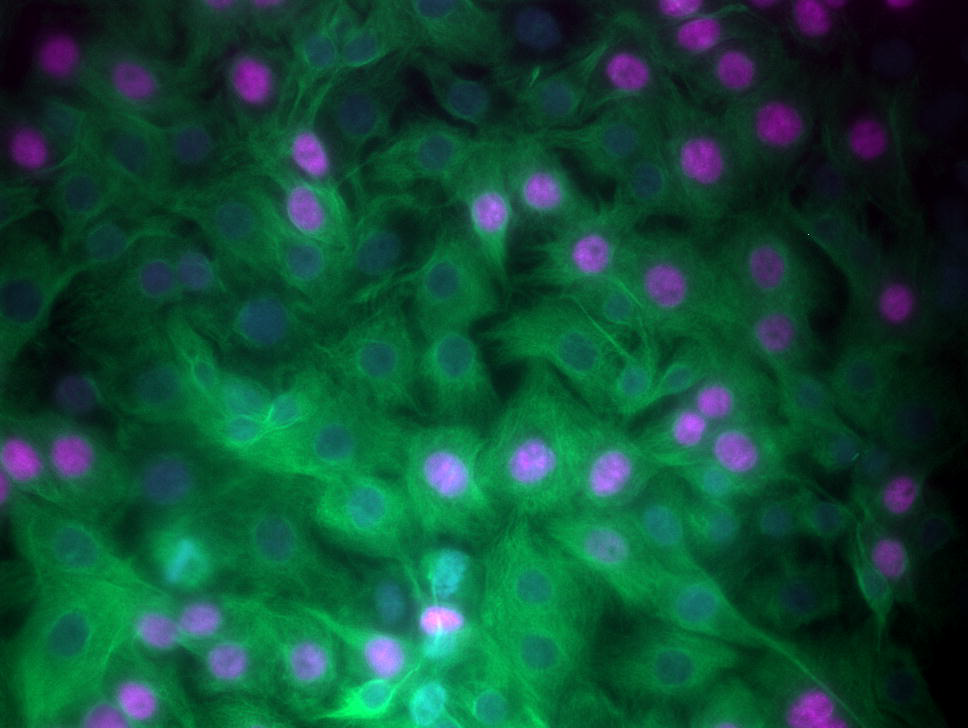

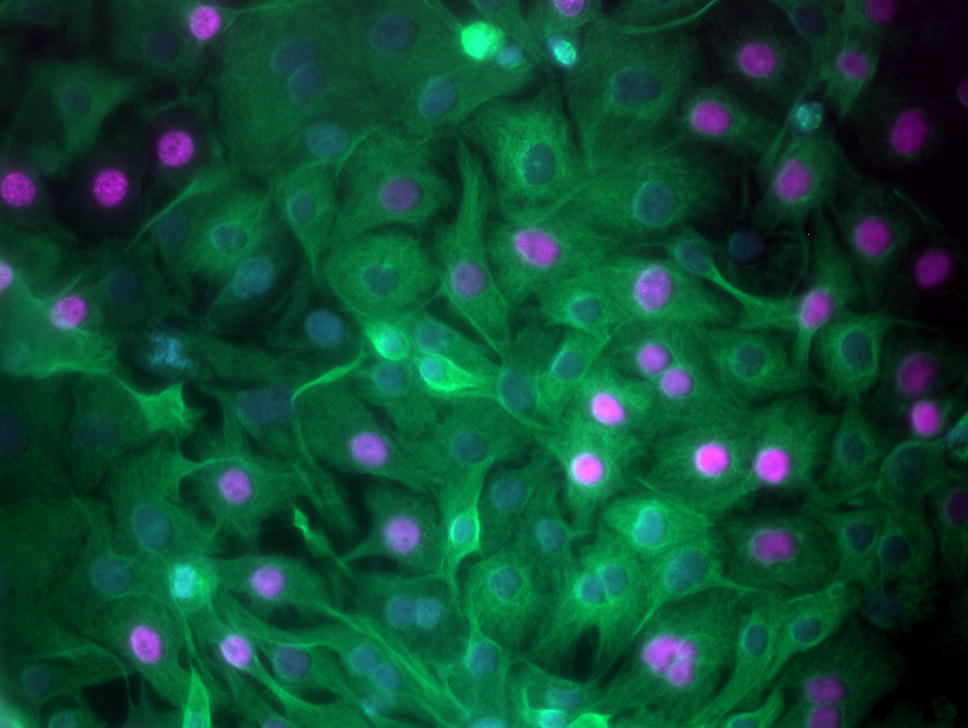

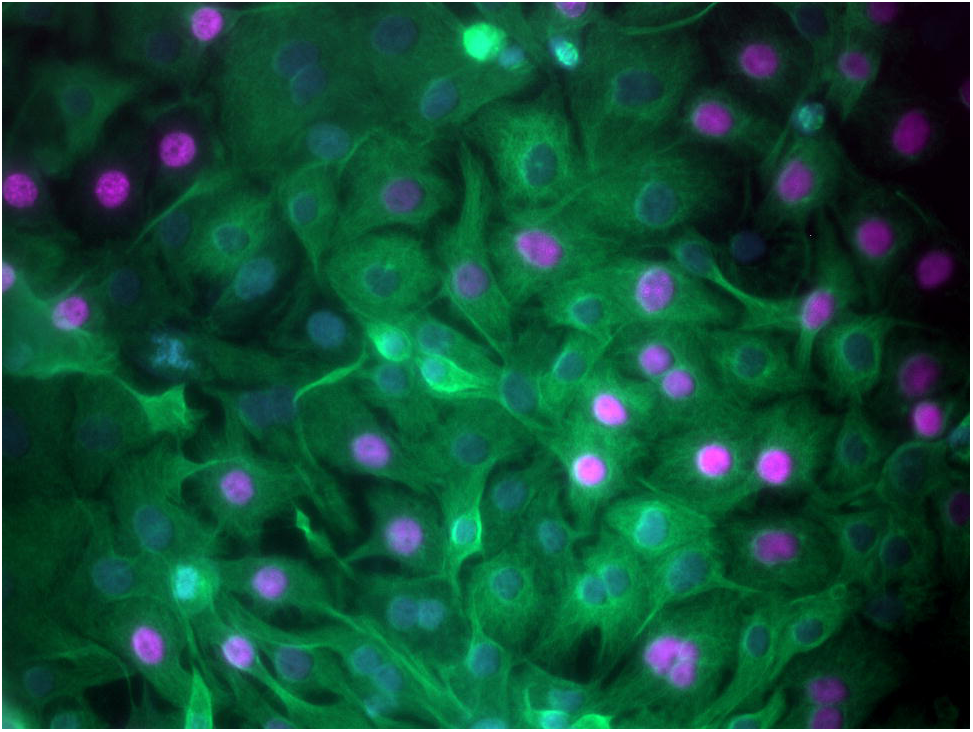

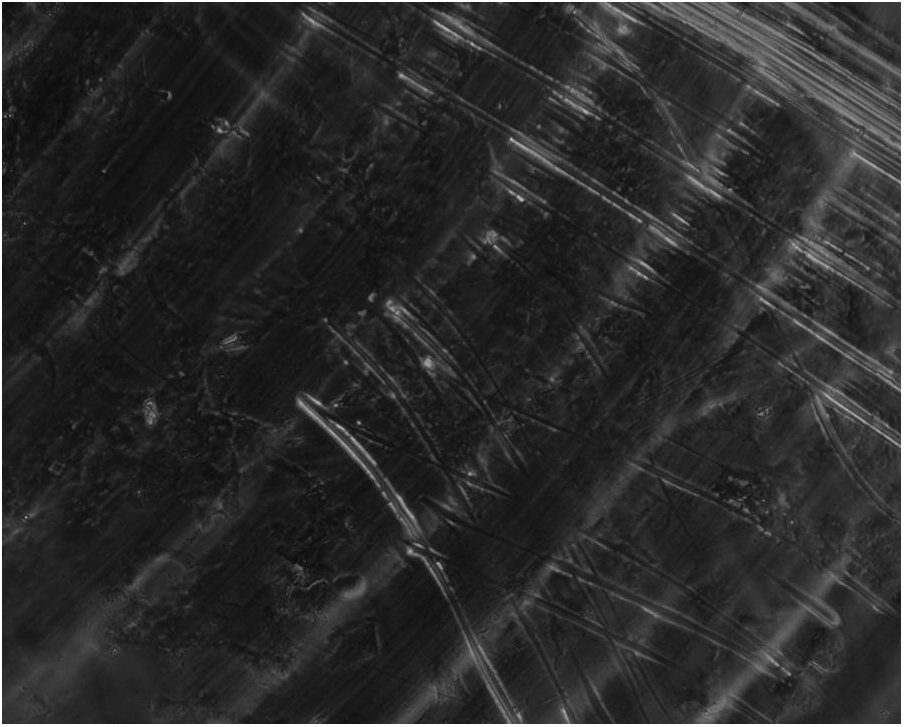

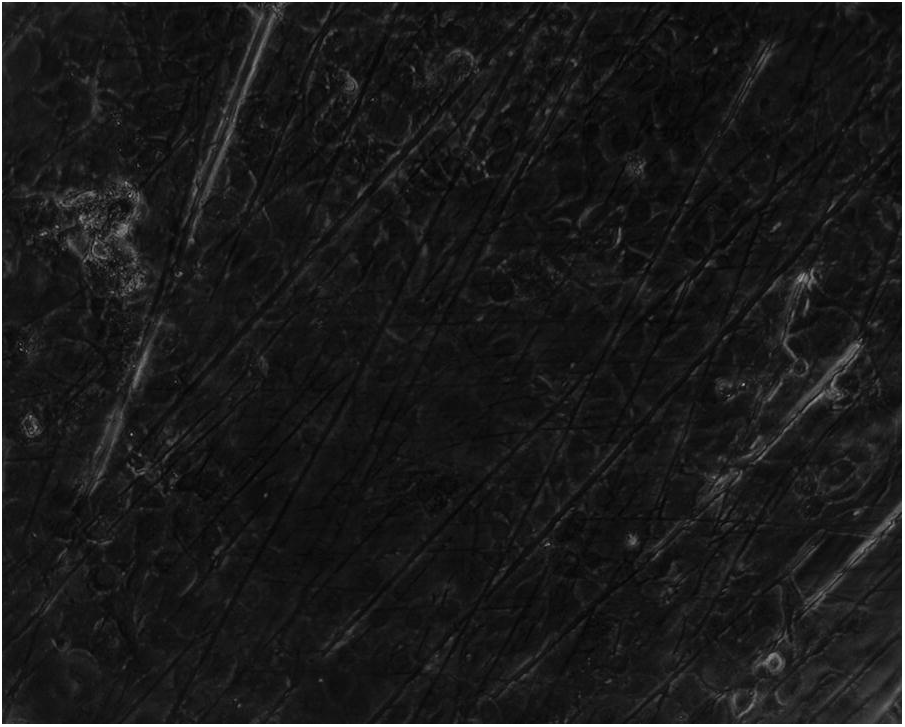

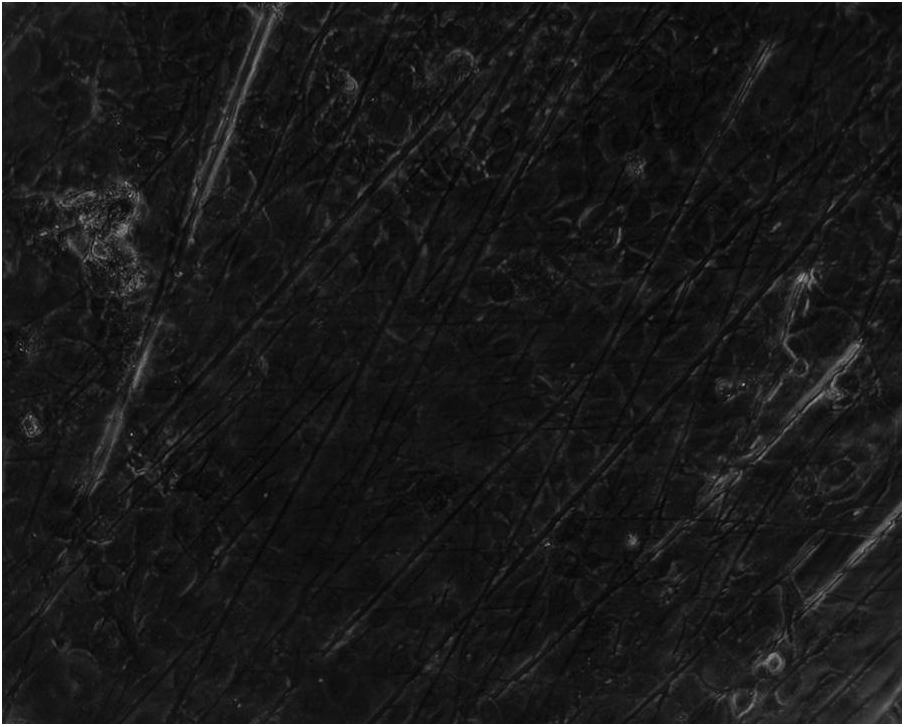

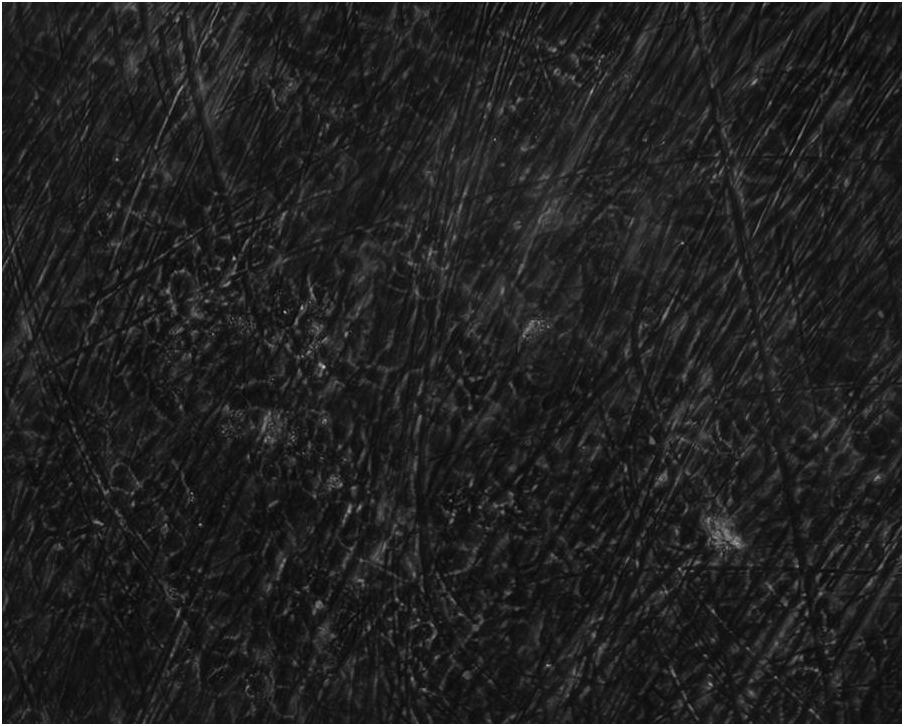

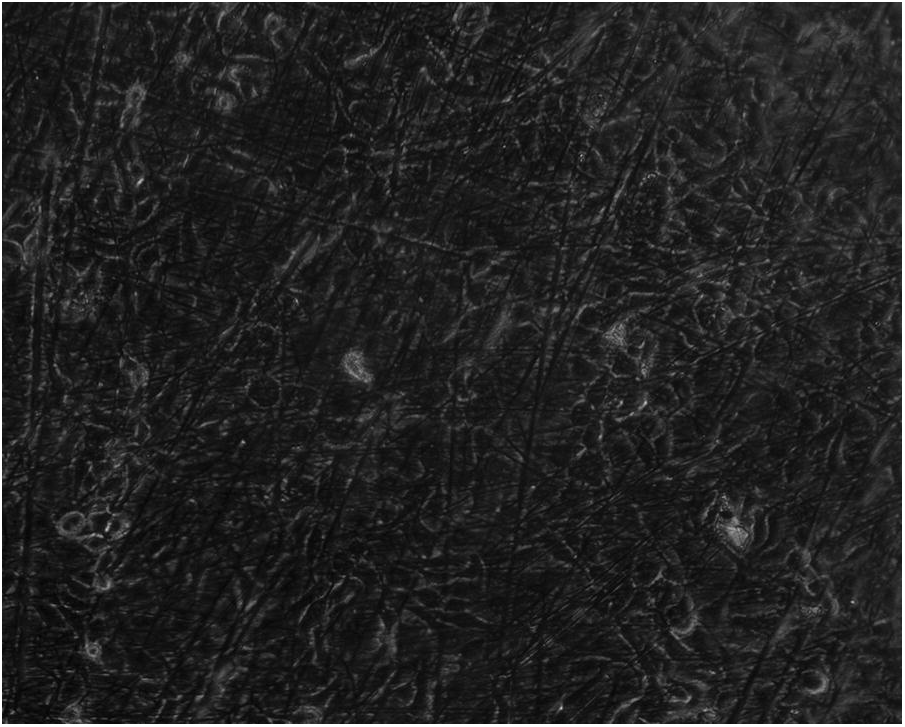

